# Differential base-sharing between humans and Neanderthals: inter-breeding or greater mutability in heterozygotes?

**DOI:** 10.1101/664581

**Authors:** William Amos

## Abstract

The idea that humans interbred with other Hominins, most notably Neanderthals, is now accepted as fact. The finding of hybrid skeletons shows that fertile matings did occur. However, inferences about the size of the resulting legacy assume that back-mutations are rare enough to be ignored and that mutation rate does not vary. In reality, back-mutations are common, mutation rate does vary between populations and there is mounting evidence that heterozygosity and mutation rate covary. If so, the large loss of heterozygosity that occurred when humans migrated out of Africa would have reduced the mutation rate, leaving Africans to diverge faster from our common ancestor and from related lineages like Neanderthals. To test whether this idea impacts estimates of introgressed fraction, I calculated D, a measure of relative base-sharing with Neanderthals, and heterozygosity difference between all pairwise combinations of populations in the 1000 genomes Phase 3 data. D and heterozygosity difference are ubiquitously negatively correlated across all comparisons, between all regions and even between populations within each major region including Africa. In addition, the larger sample of populations in the Simons Genome Diversity project reveals a pan-Eurasian correlation between Neanderthal and Denisovan fraction. These correlations challenge a simple hybridisation model but do seem consistent with a model where more heterozygous human populations tend to diverge faster from Neanderthals than populations with lower heterozygosity. Indeed, the strongest correlation between Neanderthal content and geography indicates and origin where humans likely left Africa, exactly mimicking the pattern seen for loss of heterozygosity. Such a model explains why evidence for inter-breeding is found more or less wherever archaic and human populations are compared. How much of variation in D is due to introgression and how much is due to heterozygosity-mediated variation in mutation rate remains to be determined.

**Author summary:** The idea that humans inter-bred with related lineages such as Neanderthals, leaving an appreciable legacy in modern genomes, has rapidly progressed from shocking revelation to accepted dogma. My analysis explores an alternative model in which mutation rate slowed when diversity was lost in a population bottleneck as humans moved out of Africa to colonise the world. I find that, across Eurasia, the size of inferred legacy closely matches the pattern of diversity loss but shows no relationship to where human and Neanderthal populations likely overlapped. My results do not challenge the idea that some inter-breeding occurred, but they do indicate that some, much or even most of the signal that has be attributed entirely to archaic legacies, arises from unexpected variation in mutation rate. More generally, my analysis helps explain why inter-breeding is inferred almost wherever tests are conducted even though most species avoid hybridisation.

## Introduction

It is widely accepted that humans inter-bred with related lineages such as Neanderthals and Denisovans (1–5) and that hominin inter-breeding often occurred when species’ ranges overlapped (6). The discovery of convincingly hybrid skeletons (6, 7) appears to lend unquestionable support to the fact that inter-breeding did occur. Moreover, large genetic legacies have been inferred for Neanderthals, at around 2% across Eurasia (4), and at even higher levels for Denisovan introgression into Oceania (5, 8, 9). There is also evidence that archaic DNA has been selected, with apparently introgressed haplotypes found in Tibetans living at high altitude (10) and against introgressed meiotic genes (11).

The quantity of introgressed DNA present in modern genomes is usually assessed using one of a number of versions of the so-called ABBA-BABA test and its associated D statistic (2, 12, 13) and F statistic (14) derivatives. In its basic form, the test focused on four aligned sequences, two from different human populations, one African and one European, the Neanderthal and an out-group, the chimpanzee (2). Sites are counted where the chimpanzee base, always called ‘A’ and Neanderthal base, always ‘B’, differ and where the two humans also carry one A and one B. Consequently, there are two possible states, called ‘ABBA’ and ‘BABA’. D(*San, French, Neanderthal, chimpanzee*) is calculated as the normalised difference in numbers, (nABBA – nBABA) / (nABBA + nBABA), where *n* indicates ‘number of’ counted across the genome, and was found to be around 5%.

The ABBA-BABA test and its derivatives (9, 15) explicitly assume that back-mutations are so rare they can be ignored and that mutation rate is constant (2, 16). The same two assumptions also apply implicitly in any analyses based on infinite sites models or where it is necessary to define ‘derived’ and ‘ancestral’ variants (17), since the concept of derived becomes unclear at sites carrying back-mutations. The assumption that back-mutations are rare appears to be rooted in calculation based on the genome-wide average mutation rate. However, mutation rate varies greatly across the genomic regions, with hotspots and cold spots (18–20), and back-mutations could become far more likely in hotspots. In reality, over a quarter of a million sites across the human genome in the 1000 genomes dataset carry three alleles, implying well over half a million sites with back-mutations (Amos, submitted).

The assumption of a constant mutation rate is also open to challenge. Mutation rates vary between many species, including primates (21). One mechanism that might cause variation in mutation rate between populations is heterozygote instability. HI (22, 23). The HI hypothesis is based on the observation that, during meiosis, in the synaptanemal complex, extensive regions of heteroduplex DNA are formed in which heterozygous sites become mismatches (24, 25). Such mismatches are recognised and ‘repaired’ by gene conversion-like events (26, 27), a process that has been documented in great detail in yeast and likely operate across all diploid eukaryotes. Consequently, DNA surrounding heterozygous sites will tend to experience an extra round of DNA replication in which additional mutations can occur (23, 28), implying that heterozygosity and mutation rate may be positively correlated.

Evidence for HI operating in humans comes mainly from fast-evolving microsatellites, where mutation rate covaries with both heterozygosity (29) and modern population size (30). Substitution rates also correlate with genome-wide heterozygosity and with the amount of heterozygosity lost ‘out of Africa’ (31, 32). Over all sites, human mutation rate appears marginally lower in Africans than non-Africans (33) but is substantially higher in Africans when only rare variants that likely arose after humans migrated out of Africa are considered (31). Direct sequence data from plants and insect also reveal a mutagenic effect of heterozygosity (32). Finally, different three-base combinations show markedly different mutation probabilities in different human populations (34, 35) an observation that has been extended more generally to sequence context (36). That the types of mutations differ so strongly between human populations lends credence to the idea that the overall rate might also vary.

If HI operates in humans, the large loss of heterozygosity humans experienced during the out of Africa bottleneck (37, 38) would have reduced the mutation rate in non-Africans relative to Africans. Africans would therefore have diverged more from our common ancestor and hence from related lineages than non-Africans. Such a pattern contravenes the assumptions that underpin the ABBA-BABA test and may potentially generate false signals of archaic introgression. Here I test this hypothesis by analysing the relationship between heterozygosity and D in two large genomic data sets.

## Results

### a) Simulated data

To understand the extent to which introgression and an out of African bottleneck have the potential to generate correlations between heterozygosity and D I ran a series of stochastic simulations using the coalescent simulator ms (39). I generated simulated data for two scenarios, one with no introgression and one with a level of Neanderthal introgression that was adjusted to generate realistic values of simulated D(*Africa, Europe, Neanderthal, chimpanzee*). For both scenarios, I then explored a range of ‘out of Africa’ bottleneck strengths from none through to one that erodes ~30% of the initial heterozygosity. The resulting data generate a wide range of heterozygosity differences that more than covers the full range seen in humans. In line with theory, D is large (~5%) when introgression is present and zero when introgression is absent but in neither scenario does D vary across a wide range of heterozygosity differences, generated by varying the intensity of the bottleneck. On the other hand, the level of introgression and heterozygosity in the recipient population are strongly correlated despite the modest amount of DNA this involves (Figure 1), with heterozygosity increasing in the recipient (simulated non-African) population in proportion to quantity of introgressed material.

**Figure 1.**
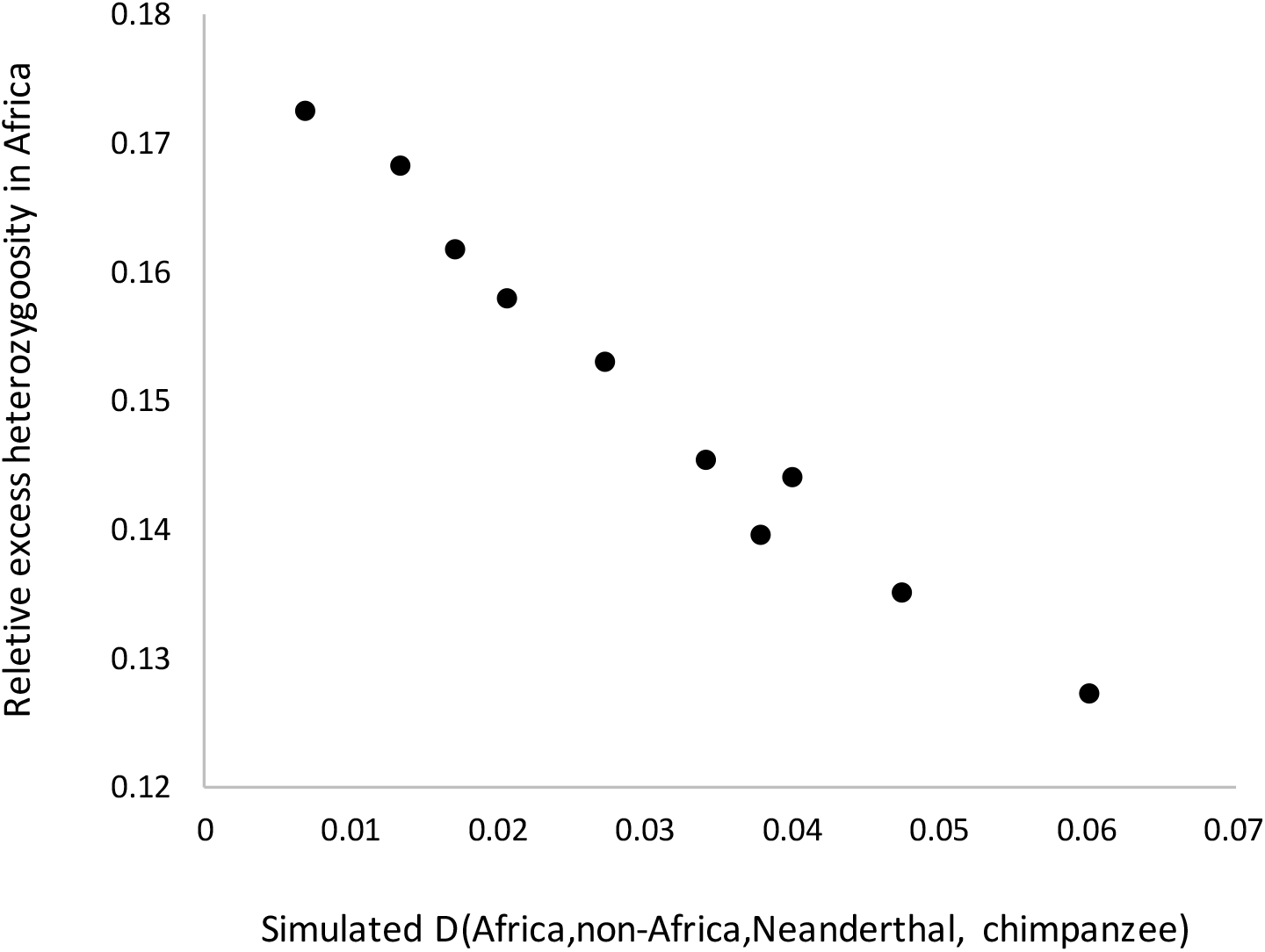
Introgression increases heterozygosity in the recipient population. Coalescent simulations were conducted using the standard scenario (see methods) which includes Neanderthal introgression, adjusted to generate D ~0.05 and a bottleneck that removes 25% of initial heterozygosity in non-Africans. From this default model, introgression was systematically varied and the resulting D values plotted against heterozygosity lost, calculated as (H_AFR_ – H_EUR_) / (H_AFR_ + H_EUR_), where H_AFR_ and H_EUR_ are heterozygosity in Africa and outside Africa respectively. Note how relative heterozygosity lost declines as the level of introgression, measured as D, increases, indicating that introgression appreciably reduces the apparent impact of the bottleneck.

### b) Relationship between D and genome-wide heterozygosity

For a broad view of the relationship between D and heterozygosity I plotted genome-wide values for all pairwise population comparisons using the 1000 Genomes Phase 3 data (Figure 2a). A strong negative relationship indicates that Neanderthals generally have a stronger affinity to the population with lower heterozygosity. However, this relationship looks to be dominated by the African – non-African comparisons, the classic pattern (2, 16) that is readily explained by an inter-breeding event soon after humans migrated out of Africa. To determine whether heterozygosity difference and D are correlated beyond African – non-African comparisons, I next plotted all pairwise comparisons between non-African populations, colour coded according to the major geographic region combinations. Overall, a strong negative relationship is again seen, with comparisons within each region-pair combination tending to form an identifiable loose cluster (Figure 2b). The American populations vary greatly in levels of European and other admixture and so comparisons in which they are involved (red, green, purple) show correspondingly large variation in heterozygosity difference.

**Figure 2.**
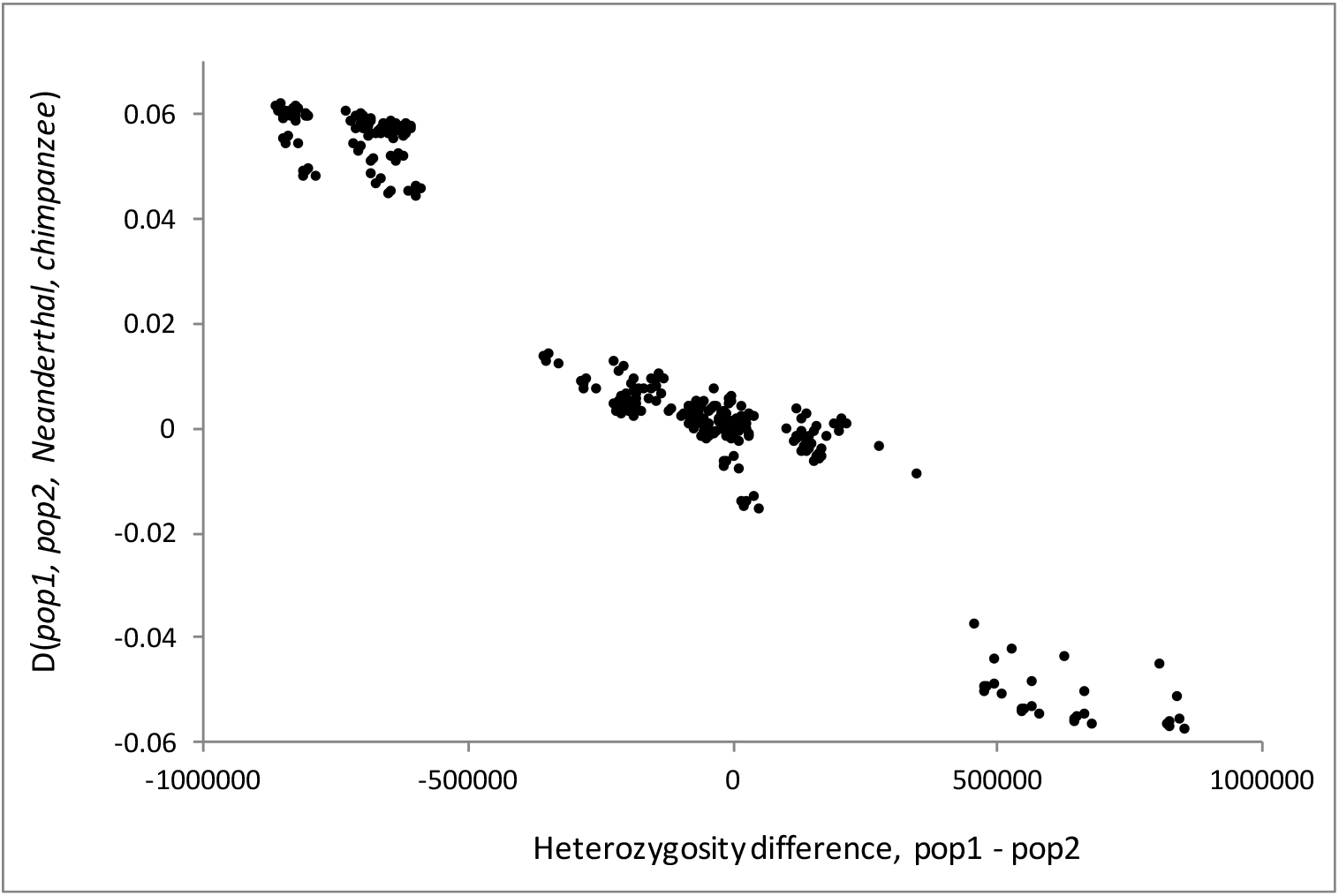
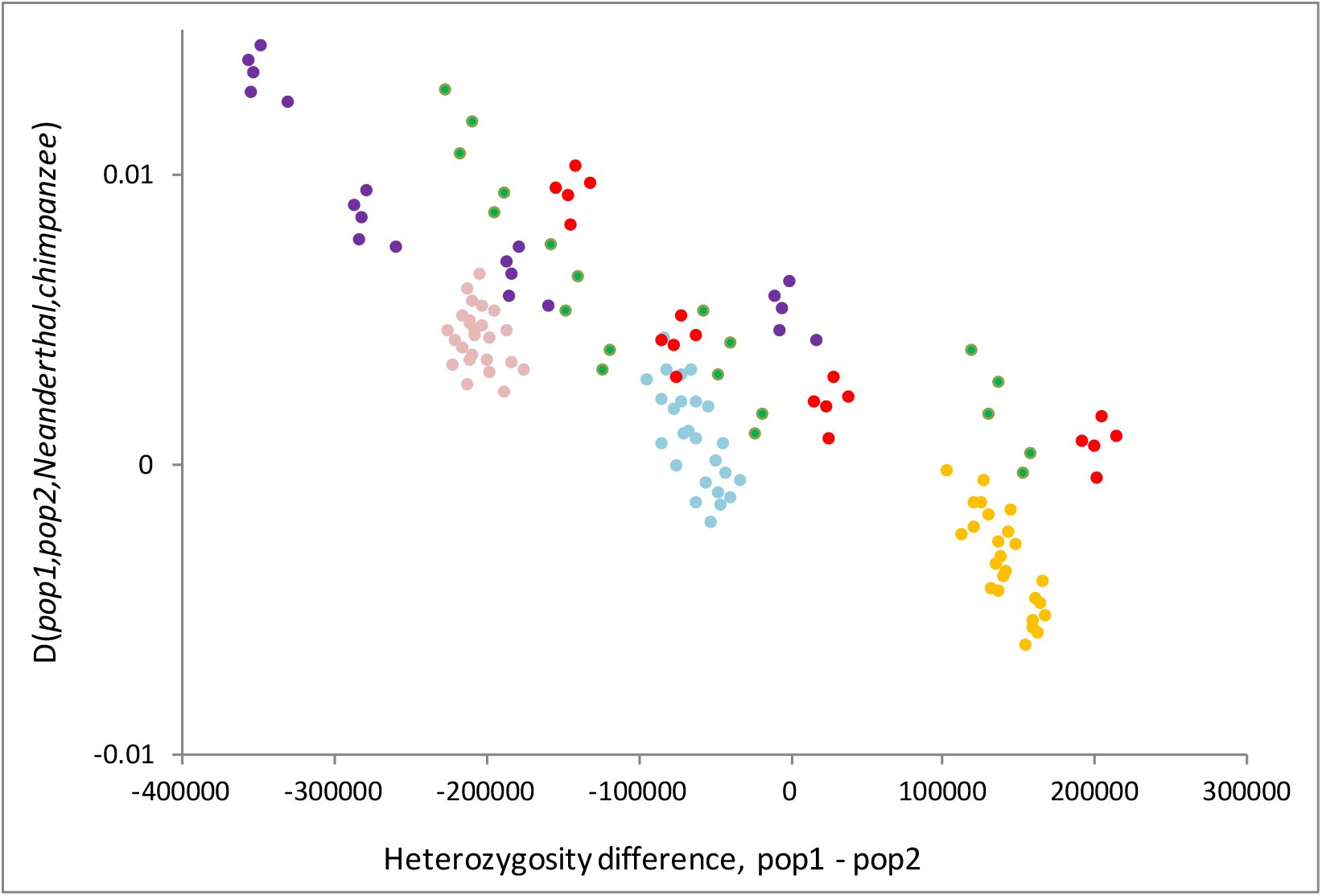
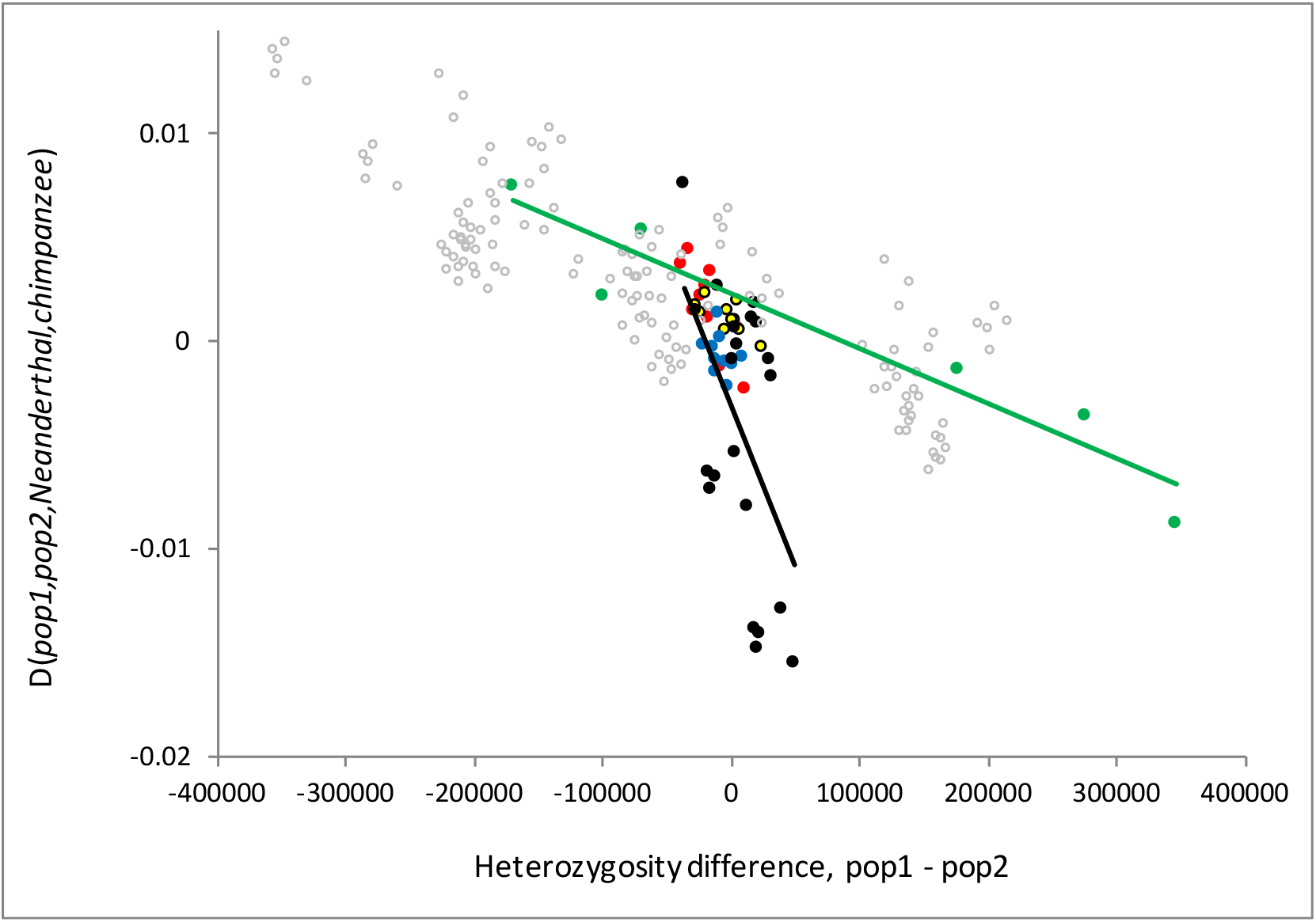
Relationship between genome-wide heterozygosity difference and D in the 1000g populations. Figure 2a, all data. Each population comparison is included once, meaning that in each comparison each population appears either as pop1 or pop2 but not both. D is calculated as (BABA-ABBA)/(BABA+ABBA): the cluster of points in the top left are all non-African – African comparisons and show positive D, indicating greater affinity between Neanderthals and non-Africans, and strongly negative heterozygosity difference, indicating higher heterozygosity in Africa. Figure 2b, between regions outside Africa: purple=EAS-AMR; green=EUR-AMR; red=CSA-AMR; pink=EAS-CSA; blue=EUR-CSA; yellow=EUR-EAS. EUR=Europe, CSA=Central Southern Asia, EAS = East Asia, AMR = America. Figure 2c, within regions: black=Africa, red=EUR, yellow=EAS, blue=CSA, green=AMR, grey circles=inter-regional combinations for comparison. Linear regressions are fitted for Africa and AMR.

I also examined comparisons *within* each of the five main geographic regions (Figure 2c). In all cases, the slope of the relationship between heterozygosity and D is negative. Two regions stand out, Africa (black) and America (green). Africa reveals a steep slope, with little variation in heterozygosity coupled with large variation in D. America shows the converse pattern, with large variation in heterozygosity but little variation in D. Despite having only four to seven populations in each group, Mantel tests reveal two significant regressions (Europe, Africa) and two borderline significant regressions (East Asia, America) (Table 1). Extending the within group analyses to between region comparisons reveals a similar pattern, with all slopes negative, many significantly so (Table 1).

**Table 1.** Mantel tests for a relationship between D and heterozygosity difference for various population combinations. All slopes are negative, regardless of whether the test is significant. EUR-EUR indicates all pairwise comparisons among European populations. EAS-CSA indicates all pairwise combinations of East Asian and Central Southern Asia populations. AMR = Africa, AFR = Africa.

| comparison | p-value |
| --- | --- |
| EUR-EUR | **0.033** |
| EAS-EAS | 0.089 |
| CSACSA | 0.421 |
| AFR-AFR | **0.007** |
| AMR-AMR | 0.069 |
| EUR-EAS | **0.037** |
| EUR-CSA | **0.022** |
| EUR-AMR | **0.008** |
| EAS-CSA | 0.245 |
| EAS-AMR | **<0.0001** |
| CSA-AMR | **<0.0001** |
| EUR-AFR | 0.059 |
| EAS-AFR | 0.051 |
| CSA-AFR | 0.088 |
| AMR-AFR | **0.03** |

### c) Introgression and geography

If heterozygosity drives mutation rate variation and this in turn drives signals interpreted as introgression, the well-documented and striking decline in heterozygosity with distance from Africa (37) should be mirrored by a general *increase* in Neanderthal legacy with distance from Africa. To test this prediction, I turned to the Simons Genome Diversity Project data (33), which comprises 300 genomes from 142 diverse populations sampled from across the world. I focused on the data contained in Supplementary Table S2 (33), which include latitude, longitude and estimates of the Neanderthal and Denisovan fractions in most samples.

Three regional groups are expected to be unusual: in Africa, there is thought to be little introgression; the American samples have variable and sometimes high European admixture; and the Oceania samples are known to have high levels of Denisovan introgression (9) (1, 8). Excluding these, leaves 198 samples from across Eurasia. Among these samples, inferred Neanderthal ancestry increases strongly with distance from Africa, taken as Addis Ababa (Figure 3, r^2^ = 0.432, N=195, P<10^−16^). To confirm that this signal is linked to the ‘out of Africa’ bottleneck, I repeated the analysis for each of a large number of evenly-spaced locations across Africa and Eurasia. For each location, I calculated a regression of Neanderthal fraction on land-only distance. In a way that precisely mirrors the relationship between heterozygosity and geography and morphological variation and geography (40), the strongest regressions between distance and Neanderthal fraction are obtained for locations in North / Central Africa / the Middle East (Figure 4).

**Figure 3.**
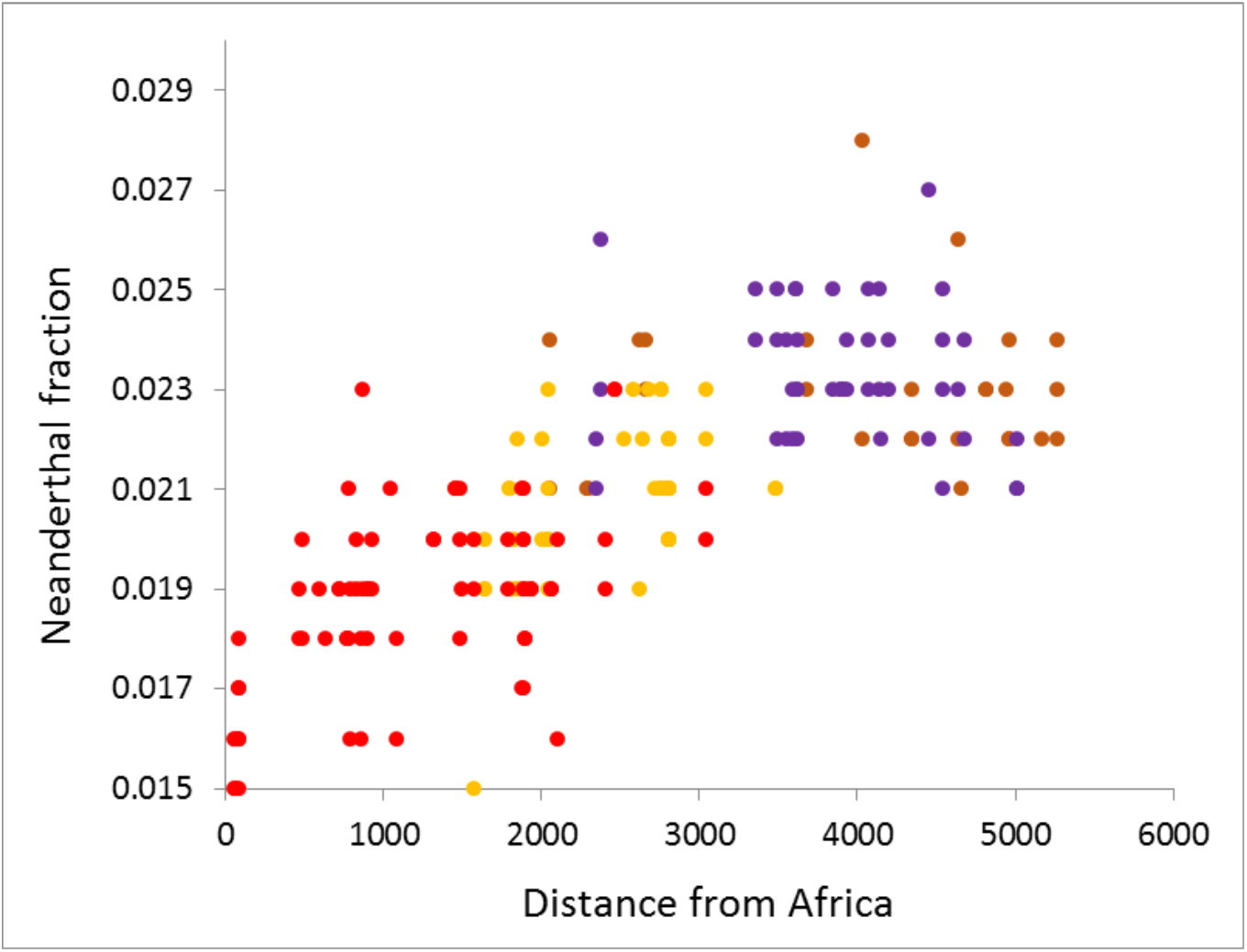
Relationship between distance from Africa and Neanderthal fraction. Values for Neanderthal fraction are taken directly from Mallick et al., Supplementary Table S2 (33). Distance is land only distance to Jordan, a point immediately outside Africa, chosen because including African data which contain many zeros would create a strong rooting at zero. Regions are: red=West Eurasia; brown=Central Asia / Siberia; purple = East Asia; yellow = South Asia.

**Figure 4.**
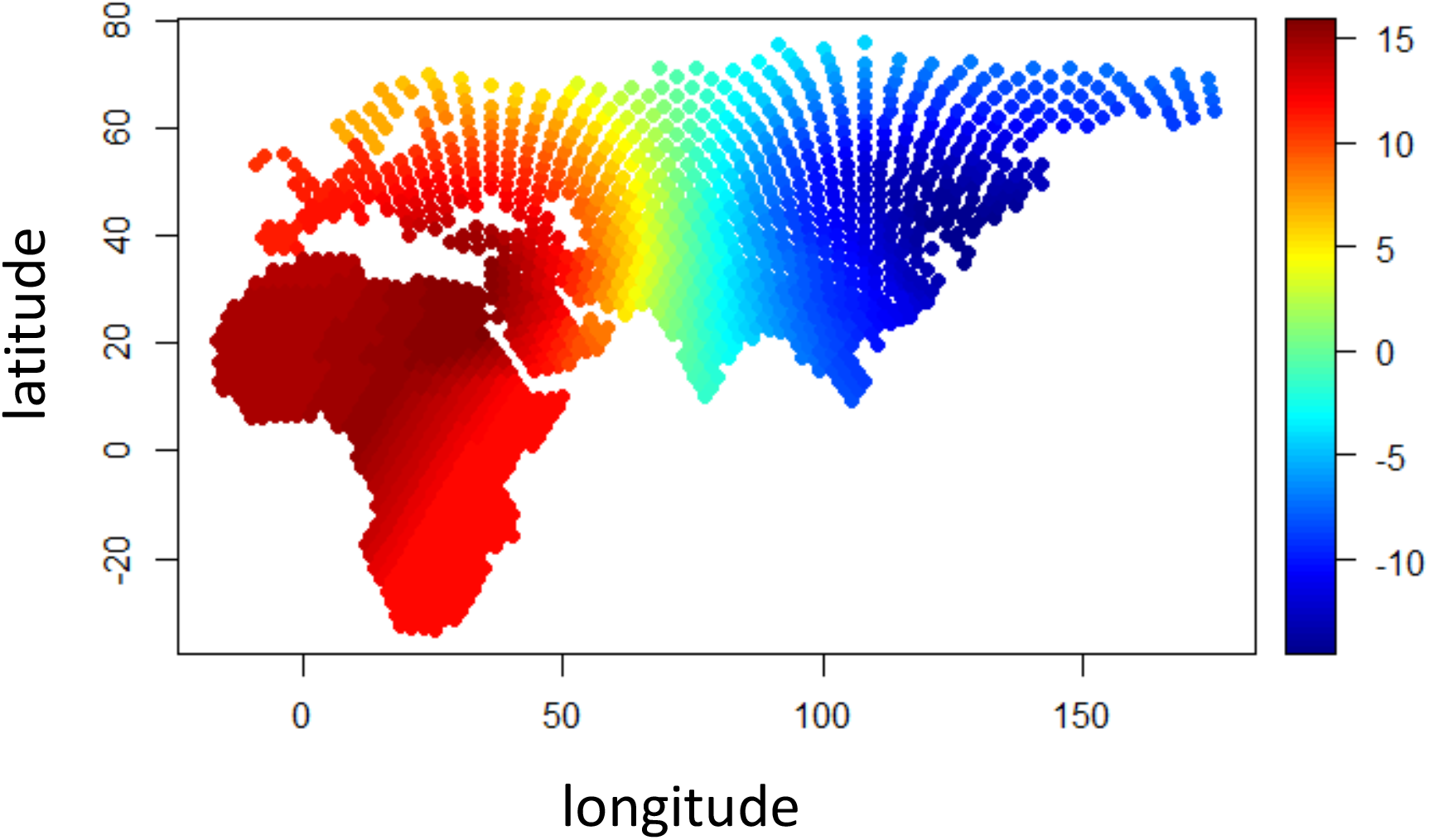
Locating the source of the strongest linear trend of Neanderthal fraction. Evenly-spaced locations were chosen across Africa and Eurasia (worldgraph.10k, geography package, R) and from each all land-only distances to the Eurasian samples in the Simons Genome Diversity Panel set were calculated and the t-value of the regression against Neanderthal fraction noted. The resulting heatmap reveals a high-point (dark red) around Northeast Egypt and a strongly positive slope indicating the Neanderthal fraction increases with distance from Africa.

Finally, Mallick et al. Supplementary Table S2 also contains estimates for the Denisovan fraction. If heterozygosity-mediated mutation slowdown contributes to signals interpreted as introgression, this should create a correlation between the Neanderthal and Denisovan fractions. This is what is found for the Eurasian data (Figure 5), a positive correlation similar in strength to the correlation between Neanderthal fraction and distance from Africa (r^2^ = 0.438, N=195, P<10^−16^). Adding in the American samples has negligible impact on r^2^ as does addition of Oceanian samples not from either Papua New Guinea of Australia, both of which have very high Denisovan content.

**Figure 5.**
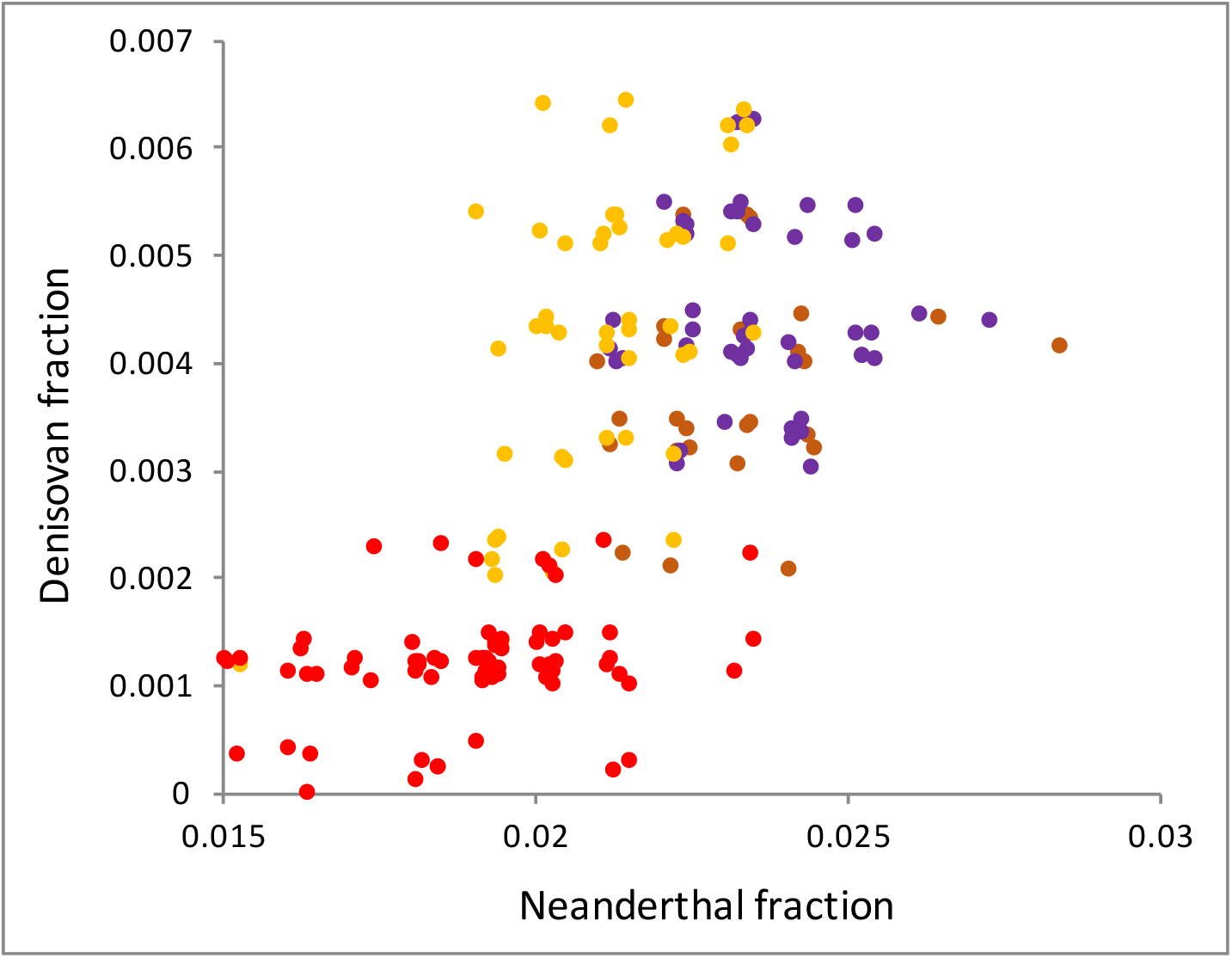
Covariation between Neanderthal and Denisovan fractions across Eurasia. Values for Neanderthal and Denisovan fractions are taken directly from Mallick et al., Supplementary Table S2 (33). Only samples from Eurasia are included. Colour coding is as for Figure 3.

## Discussion

Theoretically, the statistic often used to estimate levels of introgression, D, should be unaffected by demographic events like bottlenecks (12), even when these cause large changes in heterozygosity. At the same time, introgressed fragments will tend to increase heterozygosity such that variation in D will tend to drive variation in heterozygosity. Thus, there should be an asymmetrical relationship where variation in heterozygosity does not impact D but variation in D can cause variation in heterozygosity. This expectation is confirmed by stochastic simulations. Crucially, when variation in D drives variation in heterozygosity, the greater the level of introgression, the higher the heterozygosity of the recipient population. This is the exact opposite of what is observed in real data, where *low* heterozygosity is associated with *higher* inferred introgression.

It might be expected that geographic variation in D will reflect where Neanderthal population density was greatest, but this does not seem to be the case. Neanderthal populations appear to have been focused around South Central Europe, extending into Southwest Asia but with little or no indication of presence in North Russia, Siberia, South or East Asia or beyond (41, 42). This appears to be the exact opposite of the implied genetic legacy, which persistently increases with distance from Africa and peaks in regions where evidence of Neanderthal occupation is currently lacking. Naturally, any inter-breeding occurred a long time ago and may have pre-dated key human dispersal events such that a legacy was carried well outside regions where the two species overlap. Indeed, analysis of ancient genomes suggest a complicated history of Europeans that might help account for why the Neanderthal is higher in the East (43–45). It is also possible that the Neanderthal range is larger than current evidence suggests. Nonetheless, inferred modern legacies and what we understand about the likely Neanderthal range remain poorly matched.

A further feature of the data that sits uncomfortably with the inter-breeding hypothesis is the way the correlation between heterozygosity difference and D appears to be ubiquitous, visible inside and outside Africa and within and between regions across Eurasia. If D is driven primarily by introgression, a link to heterozygosity difference could arise either directly, through introgression (or its consequences) changing heterozygosity, or indirectly through a correlation with some other process. The direct link fails because it predicts the opposite signed slope to the one observed. An indirect link also appears unlikely because it requires humans to carry a legacy that progressively increases as they move away from where humans and Neanderthal archaeology overlap the most (41, 42). Furthermore, not only would the legacy have to increase away from Africa, but it has to increase in parallel in similar ways in South Asia, Easy Asia and Siberia.

The patterns within major geographic regions shed further light. Two regions stand out, Africa and America. In Africa, small differences in heterozygosity correlate with large changes in D while in America the converse is true. European admixture appears to play a pivotal total because the most extreme African comparisons involve Afro-Caribbeans and African Americans, while the American populations vary greatly in their levels of admixture. The African pattern is consistent both with mutation slowdown and Neanderthal introgression, since European chromosomes have little impact on heterozygosity but contribute appreciably to D regardless of the underlying mechanism. The American pattern is more informative. Under the inter-breeding hypothesis, the expectation would be that large European ancestry would increase both heterozygosity and apparent Neanderthal ancestry but the exact opposite is observed. To reconcile the observation with inter-breeding would require the European component to dilute a much higher legacy already present in Native Americans, raising questions as to why and how they carry so much more Neanderthal DNA than West / Central Eurasians.

When considering the correlations between heterozygosity difference and D it is important to remember that the overwhelming determinant of global variation in human heterozygosity is the out of Africa bottleneck. Distance from Africa explains approximately 90% of variation in heterozygosity outside Africa (37) and parallel patterns are seen for phenotypic variation (40) and gut bacterial diversity (46). This unambiguous link between demographic history and modern heterozygosity means that any trait that is strongly predicted by heterozygosity is likely driven either directly by demography or indirectly by the changes in heterozygosity that resulted. By implication, models based on selective sweeps associated with positive selection on Neanderthal alleles are unconvincing not only because they would impact just a small fraction of the genome, but also because their impact will be small relative to the loss of diversity that resulted from the bottleneck.

If inter-breeding struggles to explain the observed trends, what about the alternative hypothesis, heterozygote instability? Here, mutation rate is seen as covarying with heterozygosity such that as diversity was lost ‘out of Africa’ the mutation rate slowed causing reduced divergence both from our common ancestor and from related lineages like Neanderthals. The idea is supported by the way inferred Neanderthal fraction varies with geography, decreasing linearly with distance from Africa and revealing an African origin just like heterozygosity (40). There is also agreement on a finer scale. Thus, in the South American samples, there are large differences in heterozygosity but these are associated with small variation in D, commensurate with the short timescale of only a few centuries over which mutation rate differences driven by admixture would have had to act. Equally, the indigenous North American samples have larger, more variable and more recent admixture and reveal no trends. In a sense, these latter data help show that admixture alone is insufficient to create a consistent pattern.

The idea that heterozygosity plays some form of causative role creating phantom signals of admixture is supported by the way Neanderthal and Denisovan legacies are strongly correlated, at least across Eurasia and into America (r^2^=0.42, N=222, p<10^−16^). Such a correlation could occur if many Neanderthals and Denisovans lived side by side in time and space across Eurasia, a scenario that seems rather unlikely. A second possibility is signal leakage through shared derived alleles in the two archaics, but this would only be a strong effect if Neanderthals and Denisovans are closely related and, further, raises questions about the reliability of any one signal. The third option is that the correlated signals arise because HI impacts both signals. Where the balance lies will require further work.

In conclusion, across all population comparisons in the 1000 genomes data D, a statistic used to infer Neanderthal introgression, is predicted by difference in heterozygosity between populations. This relationship is found both inside and outside Africa, between major geographic regions and within these regions. Such ubiquity appears inexplicable by a simple introgression model, particularly since the relationship always favours populations with lower heterozygosity being closer to Neanderthals. However, the ubiquity, the direction of the relationship, the African origin and the correlation between Denisovans and Neanderthals all support the that at least part of the signals interpreted as introgression arise because the loss of diversity out of Africa slowed divergence both from our common ancestor and from related lineages.

## Methods

### Data

Data were downloaded from Phase 3 of the 1000 genomes project (47) as composite vcf files (available from ftp://ftp.1000genomes.ebi.ac.uk/vol1/ftp/release/20130502/). These comprise low coverage genome sequences for 2504 individuals drawn from 26 modern human populations spread across five geographic regions: Africa (LWK, Luhya in Webuye, Kenya; GWD, Gambian in Western Division, The Gambia; YRI, Yoruba in Ibadan, Nigeria; ESN, Esan in Nigeria; MSL, Mende in Sierra Leone; **ACB**, African Caribbean in Barbados; **ASW**, African Ancestry in Southwest US), Europe (GBR, British from England and Scotland; FIN, Finnish in Finland; **CEU**, Utah Residents (CEPH) with Northern and Western Ancestry; TSI, Toscani in Italy; IBS, Iberian populations in Spain), Central Southern Asia (**GIH**, Gujarati Indian in Texas; PJL, Punjabi in Lahore, Pakistan; BEB, Bengali in Bangladesh; **STU**, Sri Lankan Tamil in the UK; **ITU**, Indian Telugu in the UK), East Asia (CHS, Han Chinese South; JPT, Japanese in Tokyo, Japan; CDX, Chinese Dai in Xishuangbanna, China; KHV, Kinh in Ho Chi Minh City, Vietnam; CHB, Han Chinese in Beijing, China) and the Americas (**PUR**, Puerto Rican from Puerto Rico; **CLM**, Colombian in Medellin, Colombia; **MXL**, Mexican ancestry in Los Angeles, California; **PEL**, Peruvian in Lima, Peru). Some populations are likely admixed due to population history or sampling in a country far from their dominant ancestry. Codes for likely admixed populations are **bolded**. Individual chromosome vcf files for the Altai genome were downloaded from http://cdna.eva.mpg.de/neandertal/altai/AltaiNeandertal/VCF/. For analyses presented here I focused only on homozygote Neanderthal bases, accepting only those with 10 or more counts where >80% were of one base. Further data were read directly from Supplementary Table S2 in Mallick et al. (33) which also gives sample locations, population groups and estimated heterozygosity, Neanderthal and Denisovan fractions.

### Data Analysis

All analyses of the 1000g data were conducted using custom scripts written in C++, available on request. Since the 1000 genomes data are low coverage and include much imputation, population allele frequencies are determined with greater reliability than individual genotypes. Consequently, D statistics were calculated probabilistically assuming random assortment. Statistical tests were conducted using R. Geographical analyses, including calculation of land-only distances, were conducted using the R package “geoGraph”.

### Simulations

Simulated data were generated using the coalescent program ms (39). The base model was coded:

~~~
./ms 202 80000 –t 10 –I 4 1 1 100 100 –ej 6 2 1 –ej 0.3 3 2 –es 0.05
4 0.966 –ej 0.0501 5 2 –ej 0.07 4 3 –en 0.0685 4 0.007
~~~

I assume a haploid population size of 10,000, a mutation rate of 10^−8^ and set theta to 10, such that each of 80,000 non-recombining fragments is 25Kb long. The hominin-chimpanzee split is taken to be 6,000,000 years ago, the Neanderthal-human split 300,000 years ago (generation length = 25 years) and the out of Africa event 70,000 years ago. With these parameter values, an introgression event of 3.4% at 50,000 years ago generates a realistic D of ~5%. A bottleneck occurs immediately post ‘out of Africa’ with parameters set to cause a realistic average loss of 25% of initial heterozygosity.

## Acknowledgements

I thank Rob Foley, Toomas Kivisild, Andrea Manica, Simon Martin and Keziah Conroy for useful discussions. This work was not funded.

